# Differential leaf flooding resilience in *Arabidopsis thaliana* is controlled by age-dependent ORESARA1 activity

**DOI:** 10.1101/2022.11.23.517613

**Authors:** Tom Rankenberg, Hans van Veen, Mastoureh Sedaghatmehr, Che-Yang Liao, Muthanna Biddanda Devaiah, Salma Balazadeh, Rashmi Sasidharan

**Affiliations:** Plant Stress Resilience, Utrecht University; Plant-Environment Signaling; Utrecht University; Max Planck Institute of Molecular Plant Physiology; Leiden University; Laboratory of Biochemistry, Wageningen University

**Keywords:** flooding, abiotic stress, hypoxia, senescence, ethylene

## Abstract

The volatile phytohormone ethylene is a major regulator of plant adaptive responses to flooding. In flooded plant tissues, it quickly increases to high concentrations due to its low solubility and diffusion rates in water. The passive, quick and consistent accumulation of ethylene in submerged plant tissues makes it a reliable cue to trigger flood-acclimative responses, including metabolic adjustments to cope with flood-induced hypoxia. However, persistent ethylene accumulation also accelerates leaf senescence. Stress-induced senescence hampers the photosynthetic capacity and stress recovery. In submerged *Arabidopsis* shoots, senescence follows a strict age-dependent pattern starting with the older leaves. Although mechanisms underlying ethylene-mediated senescence have been uncovered, it is unclear how submerged plants avoid an indiscriminate breakdown of leaves despite high systemic ethylene accumulation. Here we demonstrate in *Arabidopsis* plants that even though submergence triggers a leaf-age independent activation of ethylene signaling via EIN3, senescence was initiated only in the old leaves, and independent of the N-degron pathway of oxygen sensing. This EIN3 stabilization also led to the overall transcript and protein accumulation of the senescence-promoting transcription factor ORESARA1 (ORE1). ORE1 protein accumulated in both old and young leaves during submergence. However, leaf age-dependent senescence could be explained by ORE1 protein activation specifically in old leaves, independent of the previously identified age-dependent control of *ORE1* via miR164. Our results unravel a mechanism by which plants regulate the speed and pattern of senescence during environmental stresses like flooding. The age-dependent activity of ORE1 ensures that older expendable leaves are dismantled first, thus prolonging the life of younger leaves and meristematic tissues vital to whole plant survival.

**Significance statement:** Flooded plants systemically accumulate saturating concentrations of the senescence promoting volatile hormone ethylene. Yet, leaf senescence follows a strict age-dependent gradient, thus prolonging the survival of young leaves and meristematic tissue. Here we show that in flooded plants, age-independent activation of ethylene signaling via EIN3, induces the systemic accumulation of the senescence-inducing transcription factor ORE1. Premature senescence of younger tissues is prevented by the posttranslational activation of ORE1 specifically in old leaves, where it induces the transcription of senescence-associated genes. Our results highlight how a systemic stress signal (ethylene accumulation upon flooding) induces a signaling cascade that diverges in an age-dependent manner, and eventually leads to an age-dependent physiological output (leaf senescence).

## Introduction

Ethylene is a gaseous hormone that controls many aspects of plant development and is a central regulator of plant environmental stress responses (Dubois et al., 2018; Sasidharan and Voesenek, 2015). Plant endogenous ethylene concentrations increase in response to a wide variety of abiotic stresses primarily mediated by enhanced ethylene biosynthesis (Argueso et al., 2007). Subsequently, ethylene triggers the stabilization of the key transcription factor ETHYLENE-INSENSITIVE1 (EIN3) leading to downstream transcriptional cascades culminating in various stress responses (Chang et al., 2013; Binder, 2020).

Flooded plants present an exception to the stress-mediated increase in ethylene biosynthesis. At least immediately following flooding, ethylene levels rapidly increase in submerged plant tissues due to physical entrapment by the surrounding flood water. This quick increase of gaseous ethylene to physiologically saturating concentrations is a consequence of the severely limited gas diffusion underwater (Xie et al., 2015; Hartman et al., 2019; Voesenek and Sasidharan, 2013). The ethylene accumulation and consequent stabilization of EIN3 is used by plants as an early flooding signal. Ethylene is a major regulator of flood adaptive traits and influences performance during flooding in various ways. For example, in deepwater rice, ethylene signaling induces stem elongation during submergence by inducing gibberellin biosynthesis and signaling (Métraux and Kende, 1983; Hattori et al., 2009; Kuroha et al., 2018). In lowland rice, on the other hand, ethylene signaling represses shoot elongation and carbohydrate consumption via SUB1A (Xu et al., 2006; Fukao et al., 2006; Fukao and Bailey-Serres, 2008). In Arabidopsis, ethylene signaling aids transcriptional responses to flood-induced tissue hypoxia, inhibits growth and modulates damage caused by reactive oxygen species (ROS) to enhance hypoxia survival (Peng et al., 2001; Liu et al., 2022; Hartman et al., 2019; Tsai et al., 2014).

The characterization of the functions of ethylene in plant flooding responses has primarily focused on traits that aid survival. However, considering its well-established role as a positive regulator of senescence (Graham et al., 2012), ethylene accumulation likely accelerates leaf senescence during submergence. Leaf senescence is often considered a marker for flood sensitivity, as flooding-intolerant accessions of rice, Arabidopsis, maize, and *Lotus japonicus* display more severe leaf senescence during flooding and post-flooding compared to tolerant accessions (Alpuerto et al., 2016; Krishnan et al., 1999; Yeung et al., 2018; Campbell et al., 2015; Buraschi et al., 2020). Furthermore, Arabidopsis mutants with reduced senescence exhibit improved performance after submergence compared to wild-type plants (Zhang et al.; Yeung et al., 2018).

The response of plant tissues to ethylene strongly depends on the age of the tissue (Doubt, 1917; Chen et al., 2013; Ceusters and Van de Poel, 2018). Ethylene treatment induces senescence much faster in older leaves than in younger leaves (dela Fuente and Leopold, 1968; Jing et al., 2005). This ensures senescence and death occur only when a leaf has reached maturity. Some mechanisms have been identified that contribute to this age-dependent response to ethylene. As a leaf ages, *EIN3* transcription gradually increases, intensifying the strength of the response to endogenous ethylene (Li et al., 2013). EIN3 induces the transcription of the master senescence regulator *ORESARA1* (*ORE1*) a NAC transcription factor, the activity of which is controlled by the kinase CALCIUM-DEPENDENT PROTEIN KINASE1 (CPK1) (Durian et al., 2020). However in young leaves premature senescence is prevented due to the degradation of *ORE1* mRNA by the microRNA *miR164* (Kim et al., 2009). As a leaf ages, the abundance of *miR164* decreases, which leads to a gradual accumulation of *ORE1*. This gradient in *miR164* works as a buffer that prevents untimely senescence in young leaves. However, increased ethylene production in stressed plants can accelerate senescence even in young leaves.

In submerged Arabidopsis rosettes that would experience systemic accumulation of ethylene, senescence still occurs along a strict leaf age-dependent gradient. Here we investigated the underlying mechanisms of this sequential leaf death. We first established that this pattern was dependent on ethylene sensing but did not require hypoxia sensing via the N-degron pathway, which is another important signaling cascade for flood acclimation. Next, we found that ethylene signaling is activated in a leaf age-independent manner upon submergence and via EIN3, induces the accumulation of the senescence-regulating transcription factor ORESARA1 (ORE1), indicating a *miR164*-independent mechanism. Although ORE1 protein was present in old and young leaves during submergence, ORE1 activation and senescence was triggered only in old leaves due to age-dependent phosphorylation of ORE1 in these tissues, independent of *CPK1*. This age-dependent phosphorylation of ORE1 ensures that leaf senescence during flooding follows an age-dependent gradient, preventing systemic tissue degradation and prolonging shoot survival.

## Results

### Ethylene perception during submergence is systemic but mediates age-dependent leaf death

Arabidopsis plants (10 leaf stage) that were completely submerged (in the dark) for varying durations exhibited a typical age-dependent pattern of leaf death. This sequential leaf death started in the oldest leaves, progressing down the age gradient towards the youngest leaves and shoot apex, which died last (Figure 1A, supplemental video 1). Ethylene has been identified as an important regulator of both flooding responses and age-dependent stress responses (Sasidharan and Voesenek, 2015; Rankenberg et al., 2021; Ceusters and Van de Poel, 2018). It has previously been established that ethylene accumulates quickly in flooded tissues (Sasidharan and Voesenek, 2015) resulting in rapid stabilization of EIN3 (Xie et al., 2015; Hartman et al., 2019). EIN3 is a transcription factor that is a key regulator of downstream transcriptional responses to ethylene (Chao et al., 1997; Chang et al., 2013).

**Figure 1.**
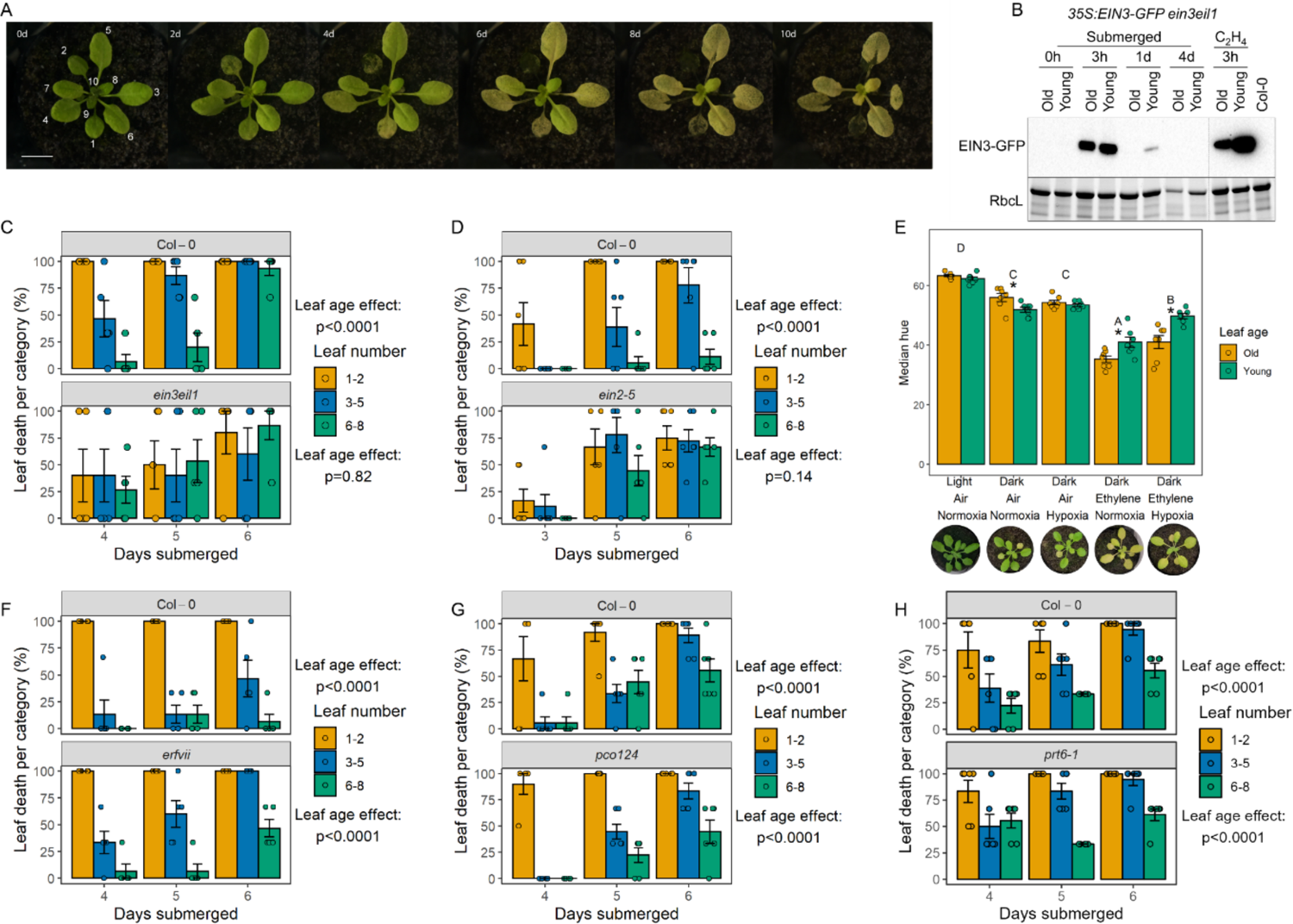
Ethylene perception during submergence is systemic but mediates age dependent leaf death. A) Arabidopsis wild type (accession Col-0) plants submerged in darkness show age-dependent leaf death. Images are representative plants submerged for the duration indicated in the top-left of each image. Numbers in the first image indicate leaf numbers. Scale bar indicates 1cm. B) Immunoblot analyses showing EIN3-GFP accumulates within 3h of submergence or ethylene treatment in both old (leaf 3) and young (leaf 7) leaves of transgenic *35S:EIN3-GFP ein3eil1* plants. Samples were run on the same gel; vertical line indicates where samples were cropped out. The large subunit of Rubisco (RbcL) served as a loading control. C) and D) Quantification of leaf death across three age-categories of an Arabidopsis rosette. Age categories are indicated by leaf number as in Figure 1A. Age-dependent leaf death observed in wild-type plants is lost in ethylene-insensitive *ein3eil1* and *ein2-5* mutants. P-values indicate the effect of leaf age on the proportion of dead leaves per genotype, determined by a two-way ANOVA (leaf age * submergence duration). n=5-6 plants per timepoint. D) The effect of different flooding cues on leaf yellowing in wild-type Arabidopsis plants. Yellowing is indicated by the median hue of old (leaf 3) and young (leaf 7) leaves, after four days of exposure to each treatment. Ethylene induces age-dependent leaf yellowing and this process is slowed down by hypoxia. Images below each bar show representative plants from each treatment. Asterisks indicate differences between old (leaf 3) and young (leaf 7) leaves (paired t-test), different letters indicate significant differences between treatments (two-way ANOVA + Tukey’s post-hoc test). n=7 plants per treatment. F - H) Age-dependent leaf death is not lost in *pco124*, *erfVII, and prt6-1* mutants, which have impaired oxygen sensing. P-values indicate the effect of leaf age on the proportion of dead leaves per genotype, determined by a two-way ANOVA (leaf age * submergence duration). n=5-6 plants per timepoint.

However, considering that the leaf death was not uniformly triggered across the submerged Arabidopsis rosettes, we wanted to establish whether ethylene signaling was indeed systemic. For this the levels of EIN3 protein were monitored in old and young leaves. Submergence enhanced EIN3 levels in both old and young leaves, already within a few hours consistent with the expected rapid accumulation of ethylene (Figure 1B). Interestingly, while EIN3 was stabilized rapidly following submergence, levels decreased thereafter during the first 24h of submergence. This is in agreement with previous observations of EIN3 as a hit-and-run transcription factor, binding briefly to its downstream targets, after which their transcription is maintained by other regulators (Alvarez et al., 2021; Chang et al., 2013). After establishing that ethylene signaling was activated systemically in flooded Arabidopsis plants, we investigated the role of ethylene in the observed leaf-age dependent senescence gradient. Age-dependent leaf death was quantified by dividing leaves into three categories based on their order of emergence: leaves 1 and 2, 3 to 5, and 6 to 8. Leaves were scored as dead when more than half of their blade area had desiccated after three days of post-submergence recovery and the proportion of dead leaves per category was calculated for each plant. This confirmed a significant leaf-age effect in wild-type plants (Figure 1C). However, this was lost in the ethylene-insensitive *ein3eil1* and *ein2-5* mutants (Figure 1C, 1D, S1B), indicating the involvement of ethylene signaling. We observed some variation between experiments in the speed at which leaves of submerged plants died. However, in all experiments the gradient in leaf death with age was consistently observed in plants that could respond to ethylene. In addition to ethylene, submergence especially during a light-limited flooding event also causes a significant decline in tissue oxygen levels (Vashisht et al., 2011; Sasidharan et al., 2018). To further probe the relative importance of ethylene in regulating the observed pattern of leaf death during submergence we exposed plants to combinations of the main submergence signals – ethylene, darkness, and hypoxia. The median hue of representative old (leaf 3) and young (leaf 7) leaves was used to quantify yellowing. The combination of ethylene and darkness induced age-dependent leaf yellowing, adding hypoxia to this combination ameliorated it (Figure 1E). However, hypoxia or darkness alone failed to trigger sequential leaf yellowing. Considering its established importance as a flooding stress cue we tested whether submergence-induced sequential leaf death requires hypoxia sensing. For this we used the Arabidopsis mutants *erfVII*, *pco124*, and *prt6-1* lacking important components of the plant oxygen sensing machinery (Abbas et al., 2015; Masson et al., 2019). In all these mutants, submergence still triggered the sequential leaf death pattern as observed in the wild type, suggesting that this response does not require oxygen sensing via the N-degron pathway (Figure 1F-H, S1B).

In conclusion, we confirmed that submergence-mediated sequential leaf death requires ethylene signaling. And, despite systemic activation of ethylene signaling in submerged rosettes, leaf death occurred in a more localized, defined pattern. The mechanisms underlying this observation were of interest to probe further.

### Submergence-induced senescence is primarily controlled by the ethylene-responsive NAC-domain transcription factor ORESARA1

The regulatory networks underpinning ethylene-mediated chlorophyll degradation leading to leaf senescence and death are well-established (Woo et al., 2019). Of relevance for submergence-induced senescence is the activation by ethylene of the NAC domain transcription factor ORESARA1 (ORE1) (Qiu et al., 2015; Yeung et al., 2018). ORE1 is a positive regulator of leaf senescence. Since the functioning of the EIN3-ORE1 regulon during leaf senescence is well-established, we used this as a system to investigate how ethylene-mediated leaf senescence is coordinated in an age-dependent manner during submergence (Kim et al., 2009; Li et al., 2013; Qiu et al., 2015).

Consistent with previous reports, ethylene-mediated senescence was reduced in *ore1-1* knockout mutants (Figure 2A; (Li et al., 2013; Qiu et al., 2015)) and ethylene exposure triggered a substantial increase in *ORE1* transcripts (Figure 2B). Interestingly, this increase was observed in both old and young leaves.

**Figure 2.**
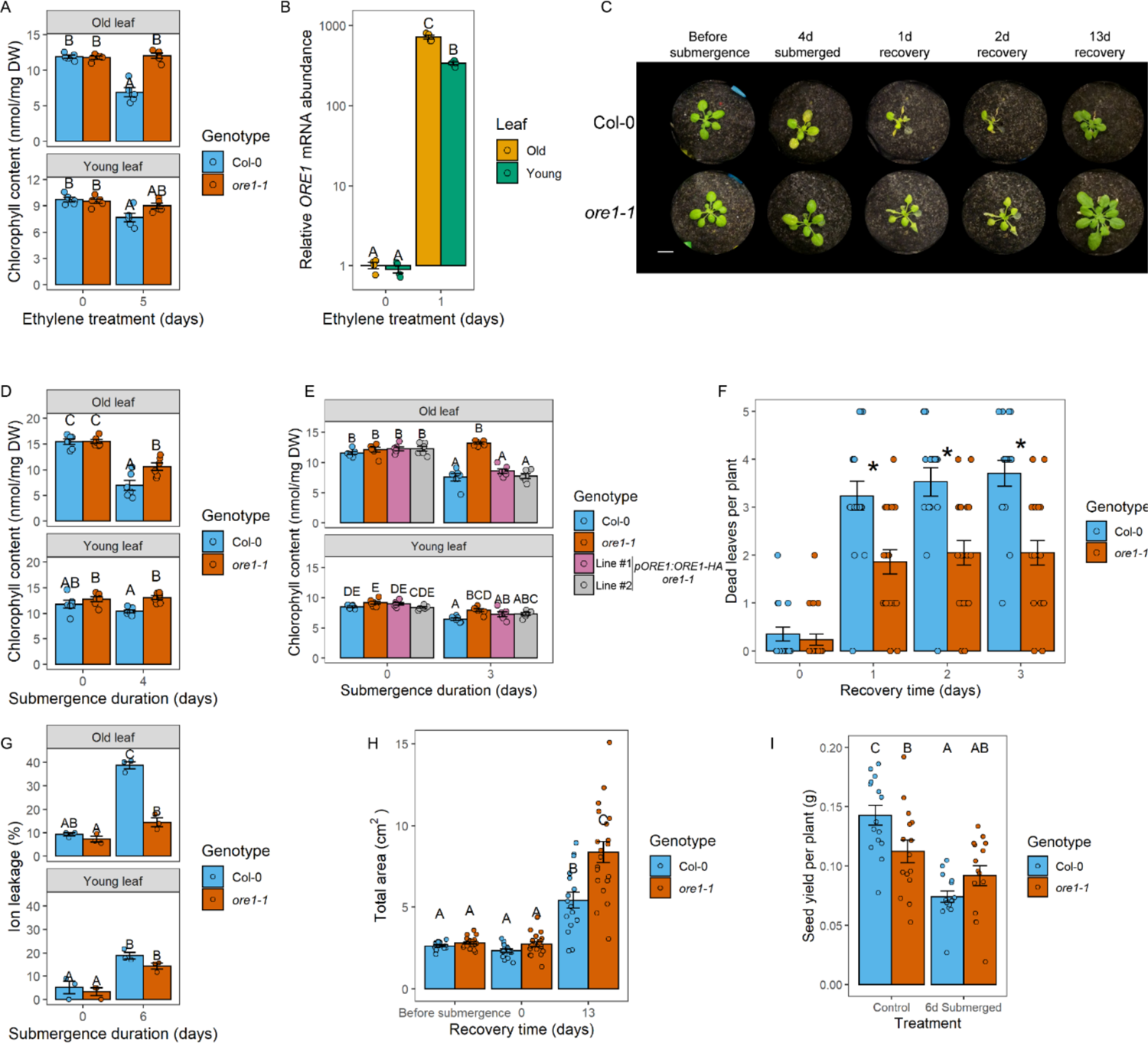
Submergence-induced senescence is primarily controlled by the ethylene responsive NAC TF ORE1. A) Chlorophyll content of old and young leaves of Col-0 and *ore1-1* plants before and after 5 days of ethylene treatment. n=5 leaves per sample. B) *ORE1* mRNA abundance increases in both old and young leaves after 1 day of ethylene treatment. n=4 leaves per sample, each consisting of 2 old or young leaves from different plants pooled together. Expression levels were normalized to those in old leaves of non-submerged plants. C) *ore1-1* mutants show reduced yellowing of old leaves after 4 days of submergence. Representative images of Col-0 and *ore1-1* plants at the indicated timepoints. Scale bar indicates 1cm. D) Chlorophyll content of old and young leaves of Col-0 and *ore1-1* before and after 4 days of submergence. n=6 leaves per sample. E) Chlorophyll content of Col-0, *ore1-1*, and two independent *pORE1:ORE1-HA ore1-1* lines before and after 3 days of submergence. n=6 leaves per sample. F) Dead leaves per Col-0 and *ore1-1* plant during recovery from 4 days of submergence. Leaves were scored as dead or alive at each of the indicated timepoints, n=17-21 plants per genotype. These same plants were phenotyped for figures S2E and S2H. G) Ion leakage of Col-0 and *ore1-1* before and after 6 days of submergence. n=3 pools of 5 old or young leaves from different plants per sample. H) Total living rosette area of Col-0 and *ore1-1* plants before and after 4 days of submergence, and after 13 days of recovery. Images of plants were categorised into dead, senescing, and healthy pixels using PlantCV. Senescing and healthy pixels were combined for each plant and converted to an area in cm^2^. n=16-20 plants per sample. I) Seed yield of Col-0 and *ore1-1* plants under control conditions and of plants that were submerged for 6 days. n=15 plants per group.

Different letters indicate significant differences between groups (two-way ANOVA + Tukey’s post-hoc test). Asterisks indicate significant differences between Col-0 and *ore1-1* per timepoint.

Next, we set out to establish that ORE1 is indeed a principal regulator of submergence-induced senescence. In accordance with its role as a positive regulator of senescence, submergence-induced senescence was significantly reduced and enhanced in *ore1* mutants and overexpressors, respectively (Figure 2C, 2D, S2A, S2B, S2C, Supplemental video 2). Moreover, the higher chlorophyll retention phenotype of *ore1-1* mutants during submergence could be reverted to wild-type by complementation with *ORE1* (ORE1 fused to an HA tag driven by its own promoter) (Figure 2E). ORE1 plays a role in dark-induced senescence of detached leaves (Kim et al., 2018). We did not detect visual signs of senescence in whole plants treated with darkness in the experimental duration used here (Figure 1D, Figure S2D) and the effect of darkness on rosette area was not different between Col-0 and *ore1-1* (Figure S2E). This shows that the submergence phenotype of *ore1-1* mutants is not merely a darkness effect. In general, higher chlorophyll maintenance in *ore1-1* mutants corresponded with a better performance during submergence relative to wild-type. This was reflected in a lower number of dead leaves and lower electrolyte leakage, although there were no significant differences in the rate of new leaf initiation immediately following desubmergence (Figure 2F, S2F, 2G, S2G). However, submerged *ore1-1* mutants also displayed a greater retention of healthy rosette area, which led to a greater rosette area after prolonged recovery (Figure S2H, 2H) and *ore1-1* seed yield was not compromised by flooding (Fig 2I). Under control conditions, *ore1-1* mutants did have a significant reduction in seed yield compared to wild-type plants. This can be attributed to delayed leaf senescence in *ore1-1* mutants. Leaf senescence plays a vital role in remobilizing nutrients from dying leaves for seed production at the end of a plant’s lifecycle (Havé et al., 2017).

Notably, we found that the phenotype of *ore1* mutants was age-dependent: the reduction in leaf senescence during flooding was most visible in old leaves (Figure 2C, S2A). Consistent with this visual observation, the decrease in chlorophyll content and cell membrane integrity was the greatest in old leaves of Col-0 plants (Figure 2D, 2G). We thus set out to investigate the regulation of ORE1 and determine how it is activated in an age-dependent manner.

### Leaf-age dependent regulation of ORE1

Submergence strongly enhanced *ORE1* transcript levels in whole rosettes and this effect was maintained also during recovery. While darkness also triggered upregulation of *ORE1,* levels quickly dropped as plants were placed back in the light (Figure 3A).

**Figure 3.**
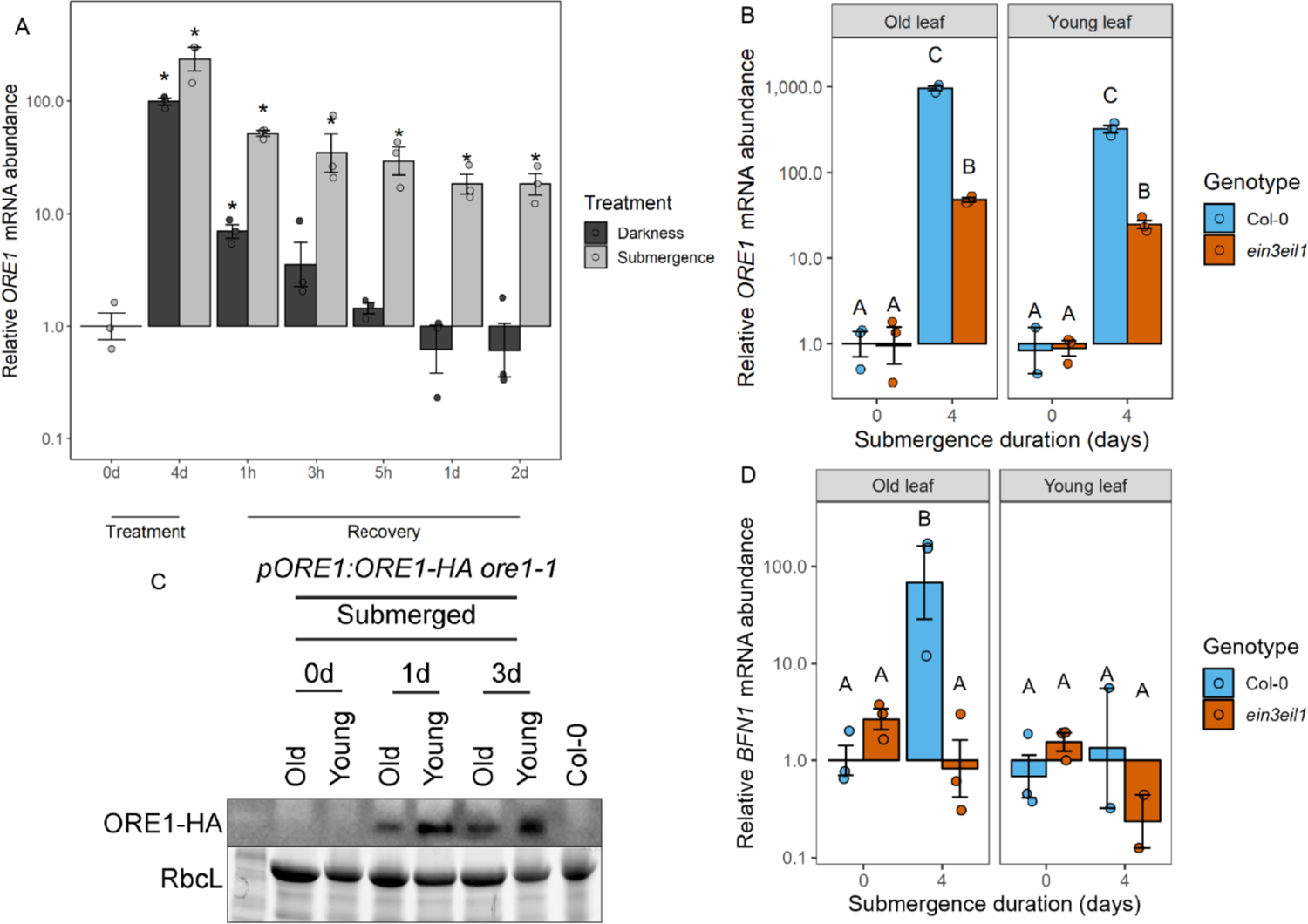
*ORE1* is induced age-independently during flooding stress. A) *ORE1* mRNA abundance in whole rosettes before and after darkness and dark submergence. Asterisks indicate significant differences compared to untreated plants (one-way ANOVA + Dunnett’s post-hoc test). Expression was normalized to that of untreated plants. n=3, each sample consists of one rosette. B) *ORE1* mRNA abundance in old and young leaves before and after four days of submergence, in Col-0 and *ein3eil1*. Different letters indicate significant differences between groups (two-way ANOVA + Tukey’s post-hoc test). Expression was normalized to that of non-submerged old leaves of Col-0. Three biological replicates were analysed, *ORE1* mRNA was not detected in one of the non-submerged Col-0 and *ein3eil1* young leaf samples; each sample consists of two leaves pooled together from different plants. C) Immunoblots showing ORE1-HA protein abundance in old and young leaves before and after one and three days of submergence, using an antibody against HA. Each *pORE1:ORE1-HA ore1-1* sample consists of five old or young leaves pooled together from different plants. Proteins of the Col-0 sample are extracted from one whole rosette. The large subunit of Rubisco (RbcL) served as a loading control. D) mRNA abundance of ORE1 target gene *BFN1* in old and young leaves before and after four days of submergence, in Col-0 and *ein3eil1*. Different letters indicate significant differences between groups (two-way ANOVA + Tukey’s post-hoc test). Expression was normalized to that of non-submerged old leaves of Col-0. Three biological replicates were analysed, *BFN1* mRNA was not detected in one of the submerged Col-0 and *ein3eil1* young leaf samples; each sample consists of two leaves pooled together from different plants.

Surprisingly, submergence led to increased transcript levels of *ORE1* in both old and young leaves (Figure 3B). This was further confirmed using a transgenic line in which the 1.6kb promoter of *ORE1* was fused to a GUS enzyme. GUS staining patterns in both young and old leaves confirmed age-independent *ORE1* promoter activity during flooding (Figure S3A). Next we examined whether age-dependent differences in *ORE1* occur at the protein level. To do so, we complemented the *ore1-1* mutant line with an HA-tagged version of ORE1, driven by its native 1.6kb promoter. In the *pORE1:ORE1-HA ore1-1* lines, there was an accumulation of ORE1 protein in both old and young leaves during submergence both at 1 and 3 days of submergence (Figure 3C). We did not detect any ORE1 protein in either old or young leaves of non-submerged plants, as short-day grown Arabidopsis plants at the 10-leaf stage have not yet initiated leaf senescence of their oldest leaves. Although *ORE1* protein and mRNA accumulated in both old and young leaves during submergence, mRNA of the ORE1 target *BIFUNCTIONAL NUCLEASE1* only accumulated in old leaves (Figure 3D). *ein3eil1* mutants still exhibited a modest increase in *ORE1* mRNA levels during submergence, but this did not lead to an increase of *BFN1* mRNA levels (Figure 3D).

Ethylene is known to enhance *ORE1* mRNA abundance via two routes – a direct transcriptional induction and via inhibition of its post-transcriptional repressor *miR164* (Kim et al., 2009). The latter mode is associated with age-dependent ethylene-induced senescence. Young leaves typically have high levels of *miR164* which decline with age. This ensures that *ORE1* mRNA is degraded when its transcription is induced by EIN3 and protects young leaves from premature senescence (Kim et al., 2009; Li et al., 2013). We did not observe significantly higher expression of *miR164b* in young leaves compared to old leaves (Figure S3B), and the similar accumulation of ORE1 protein in both old and young leaves (Figure 3C) suggests that the degradation of *ORE1* mRNA by *miR164* is not sufficient to prevent premature accumulation of ORE1 protein during submergence.

### Despite systemic ORE1 accumulation, downstream targets are activated in a leaf age-dependent manner

Although ORE1 protein accumulated in old and young leaves during submergence (Figure 3C), knocking out *ORE1* had a stronger effect on old leaves than on young leaves (Figure 2) and the ORE1 target *BFN1* was only induced in old leaves suggesting ORE1 activation only in these tissues (Figure 3D). To strengthen this evidence and get a global and unbiased overview of whether there is age-dependent activation of ORE1 targets, we carried out an mRNA Seq experiment. Old and young leaves of Col-0 and *ore1-1* were harvested before submergence, after 4 days of submergence, and after 6 hours of recovery (Figure 4A). Approximately 10 times as many differentially expressed genes (DEGs) were found between old leaves of Col-0 and *ore1-1* than between young leaves during submergence (Figure 4A). Interestingly, there were no genotype-specific DEGs when comparing the recovery timepoint to the pre-submergence timepoint. Of the 720 genotype-specific DEGs in the recovery vs submergence comparison, 428 were already differentially expressed after 4 days of submergence. This suggests that knocking out *ORE1* mostly affects the transcriptome of old leaves during submergence and not during recovery. Of the DEGs between the old leaves of Col-0 and *ore1-1* during submergence, the subset that showed a smaller increase in expression during submergence in *ore1-1* than in Col-0 contained several previously identified targets of ORE1 including (*BIFUNCTIONAL NUCLEASE1* (*BFN1*) and *NON-YELLOWING1* (*NYE1*)). Furthermore, these DEGs were enriched for ORE1 binding sites near their transcriptional start sites (Figure S4A). DEGs that did not fall within this subset did not have this enrichment, nor did non-DEGs.

**Figure 4.**
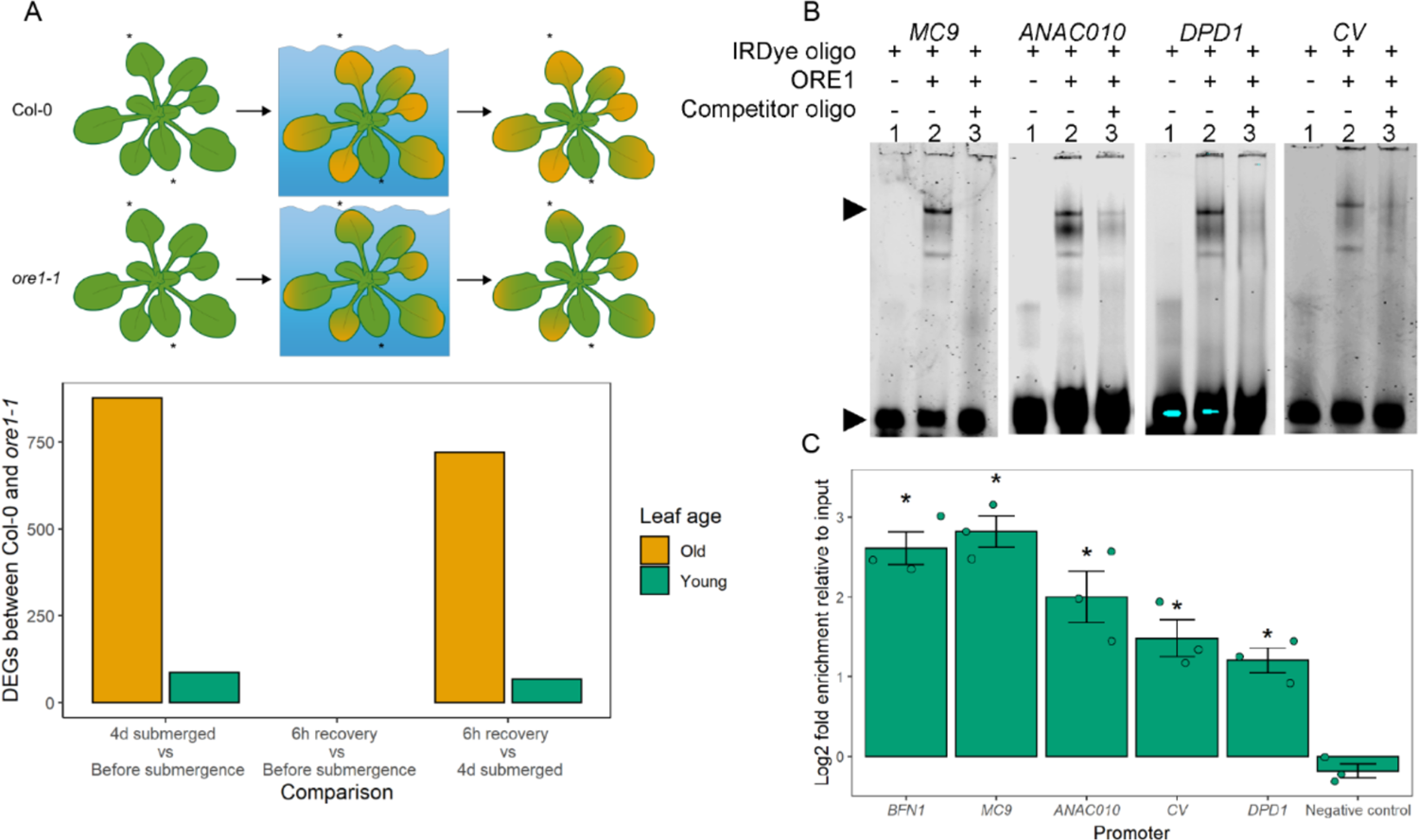
ORE1 target activation is age-independent. A) Leaf samples of Col-0 and *ore1-1* were harvested before submergence, after four days of submergence, and after 6 hours of recovery. The number of differentially expressed genes (DEGs) that show a genotype-dependent response to four days of submergence is greater in old leaves than in young leaves. None showed a genotype-dependent effect in their response to submergence followed by recovery. Most (428/720) DEGs showing a genotype-specific response to post-submergence recovery showed the opposite pattern during the submergence phase. B) Electrophoretic Mobility Shift Assay (EMSA) showing *in vitro* binding of recombinant ORE1-GST to the promoters of *MC9*, *ANAC010*, *DPD1*, and *CV*. From left to right in each image: lane 1, labeled probe (5′-DY682-labeled double-stranded oligonucleotides); lane 2, labeled probe plus ORE1-GST protein; lane 3, labeled probe, ORE1-GST protein, and competitor (unlabeled oligonucleotide containing an ORE1 binding site; 200× molar access). Arrows indicate retarded bands (bound oligo) and the non-bound DNA probes (free oligo). C) ChIP-qPCR showing *in vivo* binding of ORE1 to the promoters of *MC9*, *ANAC010*, *DPD1*, and *CV*. Asterisks indicate significant enrichment relative to the negative control (AT2G22180) (one-way ANOVA + Dunnett’s post-hoc test). Chromatin was extracted from immunoprecipitated samples from whole *pORE1:ORE1-HA* rosettes submerged for one day, n=3.

We expanded the set of known ORE1 target genes by confirming that ORE1 can bind the promoters of the protease *METACASPASE9* (*MC9*), the transcription factor *ANAC010*, the nuclease *DEFECTIVE IN POLLEN ORGANELLE DNA DEGRADATION 1* (*DPD1*), and the chloroplast-degrading protein *CHLOROPLAST VESICULATION* (*CV*) *in vitro* via electrophoretic mobility shift assay (EMSA) (Figure 4B). These new targets were selected based on their roles in senescence-related processes. *In vivo* binding of ORE1 to these promoters was confirmed via ChIP-qPCR using 1-day submerged *pORE1:ORE1-HA* plants (Figure 4C). The binding of ORE1 to all newly identified putative targets was significantly enriched, when compared to the negative control (AT4G22180) (Figure 4C). Out of a set of 15 verified ORE1 targets, from this study and others ((Rauf et al., 2013; Matallana-Ramirez et al., 2013; Qiu et al., 2015; Zhang et al., 2021); Table S1), *ORE1* disruption affected the submergence-induction of 12 (in old leaves). In young leaves, however, only 3 out of the 15 differed in their response to submergence between Col-0 and *ore1-1* (Figure S4B).

The *ORE1*-dependent response during submergence does not seem to involve hypoxia signaling, as none of the 47 out of the 51 core hypoxia genes (HRGs) (Mustroph et al., 2009) detected in our dataset were different between Col-0 and *ore1-1* in either old or young leaves (Figure S2C). The mRNA Seq data also confirmed that global ethylene signaling was induced similarly in old and young leaves during submergence, based on the similar expression of EIN3 target genes between these leaves (Figure S4D).

### Age-dependent ORE1 phosphorylation during submergence is required for downstream target activation

Although ORE1 protein accumulated to higher levels during submergence in young leaves than in old leaves, its downstream targets were mostly activated in old leaves. This indicated an age-dependent activation of ORE1 in old leaves. The transactivation ability of ORE1 was recently shown to depend on its six-fold phosphorylation (Durian et al., 2020). We thus probed this post-translational modification as a potential mechanism mediating differential ORE1 activation in our system. Protein extracts from leaves of submerged *pORE1:ORE1-HA* plants run on an SDS-PAGE gel containing 50µM PhosTag revealed a slower migration of ORE1-HA from old leaves. This suggested the presence of phosphorylated ORE1-HA in old, submerged leaves supporting our hypothesis of age-dependent ORE1 activation via phosphorylation during submergence (Figure 5A). To further validate this, we used transgenic plants overexpressing a modified ORE1 missing the region between amino acids 205 and 221 containing potential phosphorylation sites (*35S:ORE1Δ17)*. These *35S:ORE1Δ17* plants showed a phenotype intermediate between Col-0 and *ore1-1* plants under submergence stress (Figure 5B, 5C).

**Figure 5.**
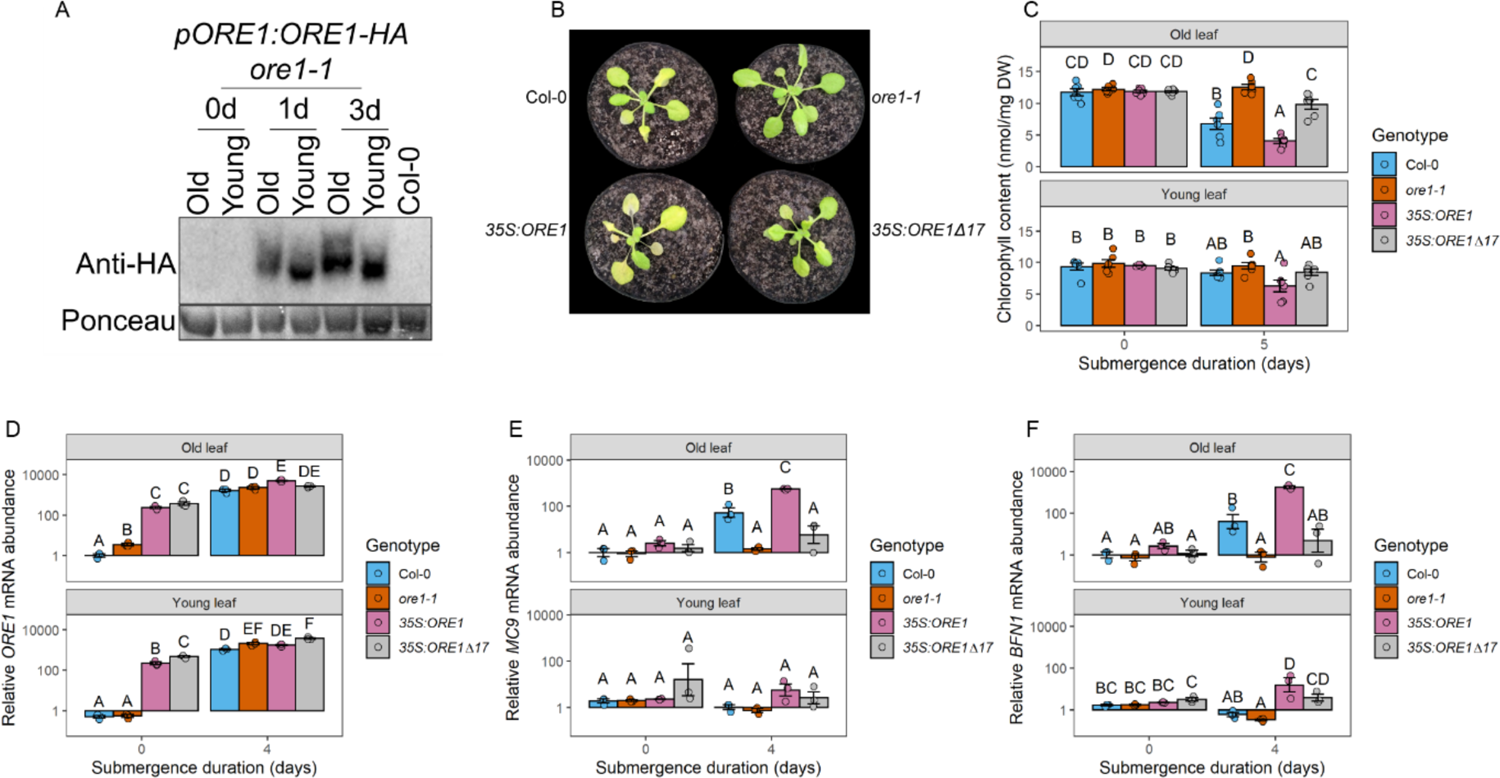
ORE1 phosphorylation during flooding is age-dependent. A) *pORE1:ORE1-HA ore1-1* protein samples from submerged old leaves move slower through a PhosTag gel than samples from young leaves, indicating age-specific phosphorylation of ORE1. Five old or young leaves were pooled together from different plants per *pORE1:ORE1-HA ore1-1* sample, the Col-0 sample was from one whole rosette. B) Representative images of Col-0, *ore1-1*, *35S:ORE1*, and *35S:ORE1Δ17* plants after 5 days of submergence followed by one day of recovery. Scale bar indicates 1cm. C) Chlorophyll content of Col-0, *ore1-1*, *35S:ORE1*, and *35S:ORE1Δ17* plants, before and immediately after 5 days of submergence. D-F) Expression of *ORE1*, *MC9*, and *BFN1* in Col-0, *ore1-1*, *35S:ORE1*, and *35S:ORE1Δ17* before and after 4 days of submergence. Expression was normalized to that of non-submerged old leaves of Col-0. Two old or young leaves from different plants were pooled together per sample. Different letters indicate significant differences between groups (two-way ANOVA + Tukey’s post-hoc test). Error bars indicate SEM.

As expected, the expression of *ORE1* was already high before submergence in *35S:ORE1* and *35S:ORE1Δ17* and was also induced in both old and young leaves of Col-0 and *ore1-1* leaves during submergence (Figure 5D). The *ore1-1* mutant is a true null mutant containing a T-DNA insertion in the last exon. The primer pair used here spans the first intron, explaining the increase in *ORE1* transcript levels in *ore1-1* (Durian et al., 2020; Balazadeh et al., 2010). Although expression of *ORE1* was high during submergence in both old and young leaves of all four genotypes used here, the downstream target genes *MC9* and *BFN1* were only induced in the old leaves of Col-0 and *35S:ORE1* (Figure 5E-F). Taken together, these results suggest that the age-dependent phosphorylation of ORE1 is required for the activation of its downstream target genes.

### Ethylene exposure is sufficient to induce age-dependent accumulation of phosphorylated ORE1

Plants with impaired ethylene signaling did not show age-dependent leaf death during submergence, and treating plants with ethylene in darkness induced age-dependent leaf death (Figure 1C-E). This could not be explained by *ORE1* transcript levels since ethylene treatment and submergence caused leaf-age independent *ORE1* induction (Figure 2B & 3B). This held true also for ORE1 protein levels (Fig 3C and Fig 6A). While the combination of ethylene and darkness was both essential and sufficient for the induction of ORE1 protein to similar levels as during submergence, this occurred in both old and young leaves (Figure 6A). However, considering that during submergence ORE1 phosphorylation and activation occurred only in old leaves, we hypothesized that ethylene might be the underlying submergence signal (Figure 5A). In accordance with this, ethylene exposure in darkness was already sufficient to induce the accumulation of age-dependently phosphorylated ORE1 protein in one day (Figure 6B). This is consistent with the previous observation that ethylene treatment, rather than hypoxia, is sufficient to induce age-dependent leaf yellowing (Figure 1D). Furthermore, treating plants with ethylene in darkness had a similar effect on the senescence phenotype of Col-0 plants as submerging them in darkness (Figure 6C, 6D). In ethylene-insensitive *ein3eil1* plants, however, this induction of senescence during dark submergence was lost. To probe this further, we induced *ORE1* expression throughout the rosette by using transgenic plants expressing *ORE1* under an estradiol-responsive promoter. Systemic *ORE1* induction led to an age-dependent induction of its target genes and age-dependent leaf yellowing (Figure S5). This suggests that flooding-induced ethylene signaling controls the systemic ORE1 accumulation but is not essential for the age-dependent activation of ORE1. The loss of age-dependent senescence observed in flooded ethylene-insensitive mutants (Figure 1), could likely be an effect of the role of ethylene in leaf development (Vandenbussche et al., 2012).

**Figure 6.**
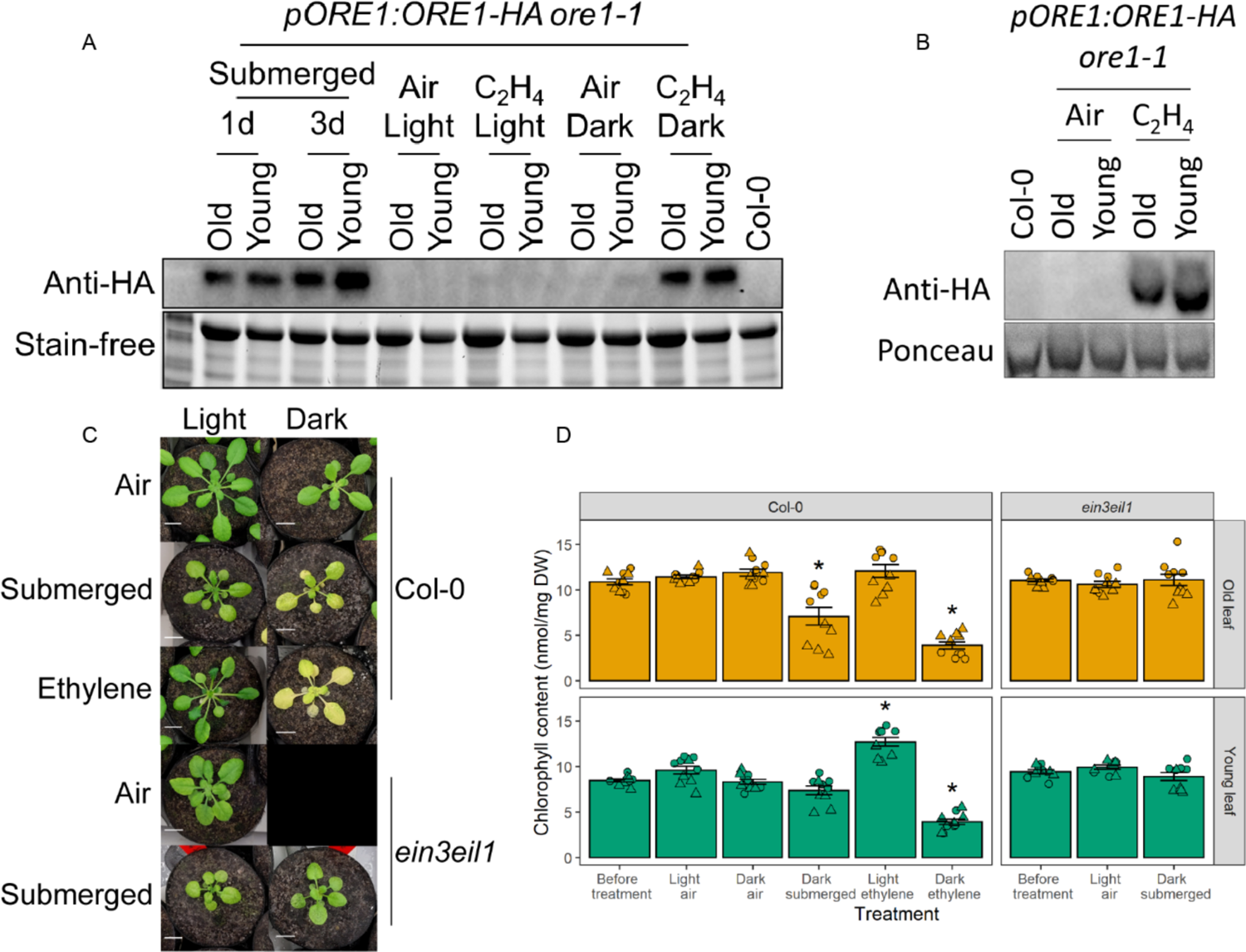
Ethylene controls leaf age-dependent ORE1 phosphorylation. A) Immunoblots showing ORE1 accumulation in old and young leaves after submergence or one day of ethylene treatment in darkness. Five old or young leaves from different plants were pooled together per *pORE1:ORE1-HA ore1-1* sample. The Col-0 sample was from one whole rosette. Stain-free imaging of the protein gel was used as a loading control. B) ORE1-HA from old leaves treated with ethylene in darkness for one day moves slower through a phostag gel than ORE1-HA from young leaves. Samples are the same as the ones run on the non-phostag gel in panel A. Ponceau staining of the large subunit of Rubisco was used as a loading control. C) Shoot phenotypes in response to submergence or ethylene in light or dark conditions. Representative images of Col-0 and *ein3eil1* plants immediately after five days of the indicated treatments are shown. Scale bars indicate 1cm. D) Chlorophyll content of old and young leaves following treatments with different combinations of ethylene and darkness. Asterisks indicate significant differences from chlorophyll levels before the treatment (One-way ANOVA + Dunnett’s test), error bars indicate SEM. n=10 per sample from two independent experiments, circles and triangles indicate experimental replicates.

ORE1 is phosphorylated by CPK1 *in vivo* (Durian et al., 2020). *CPK1* mRNA levels showed a leaf age-dependent increase in submerged plants, although the absolute changes in expression are small (Figure S6A). *CPK1* also possessed an EIN3 binding site in its promoter (Figure S6B). We thus investigated it as a candidate kinase that phosphorylates and activates ORE1 downstream of ethylene. However, *CPK1* expression did not change in response to ethylene treatment in either old or young leaves (Figure S6C). Consistent with this, chlorophyll content of *cpk1-1* mutants was not different from Col-0 after either submergence or ethylene treatment (Figure S6D-E). A comparison of Col-0, *ore1-1*, and *cpk1-1* plants revealed age-dependent leaf death in all genotypes upon submergence. This was significantly delayed in *ore1-1* compared to Col-0 but not in *cpk1-1* (Figure S6F). Thus, although ethylene exposure selectively induces senescence in old leaves via the age-dependent phosphorylation of ORE1, this does not seem to depend on *CPK1*.

Taken together, these results provide a mechanism by which plants ensure that leaf senescence follows an age-dependent gradient during flooding stress. Such a mechanism might safeguard a total overall collapse of the plant due to high ethylene accumulation during flooding. Interestingly it is still unclear what prevents ethylene-mediated activation in young leaves. Although flooding stress induces systemic ethylene signaling and ORE1 accumulation, the age-dependent phosphorylation of ORE1 ensures that it can only activate its downstream targets in older tissues (Figure 7). The accumulation of ORE1 in young leaves can prepare them to rapidly transition into senescence if the submergence duration is long enough.

**Figure 7.**
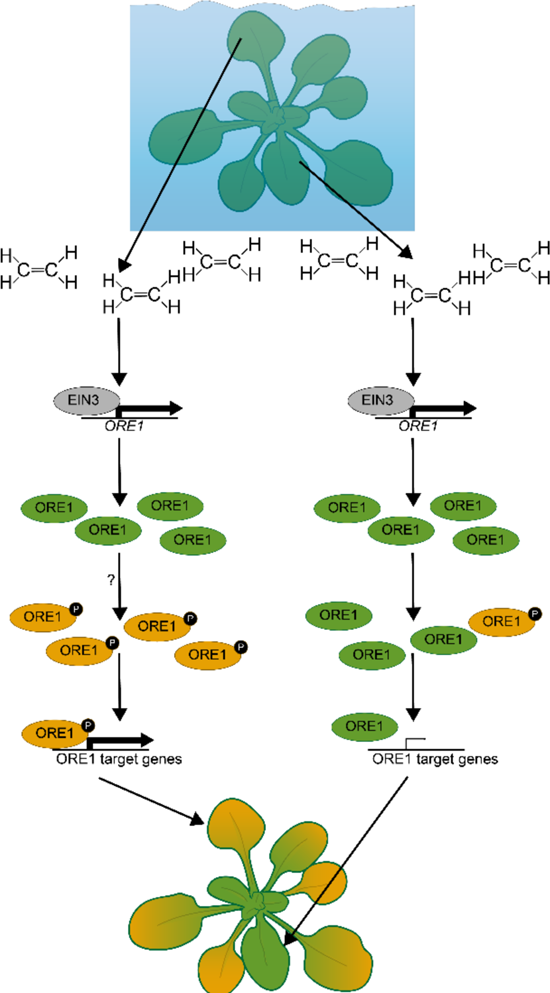
A model for ethylene-mediated sequential leaf death in flooded plants.

Upon submergence, ethylene rapidly accumulates throughout a plant. This age-independent accumulation of ethylene induces the age-independent accumulation of *ORE1* mRNA and protein via EIN3 stabilization. Leaf-age dependent senescence is triggered by ethylene via ORE1 phosphorylation and activation specifically in old leaves, via an unknown mechanism. This age-dependent phosphorylation of ORE1 ensures that it induces senescence in old leaves, which results in the oldest leaves being broken down first and the youngest leaves and meristem last.

## Discussion

Our results demonstrate a mechanism whereby plant responses to a systemic stress cue are locally determined. Submergence of Arabidopsis rosettes activates ethylene signaling in all leaves, consistent with an expected systemic accumulation of ethylene and yet initiates senescence in a specific leaf-age dependent pattern. Leaf senescence during flooding starts in the oldest leaves, but eventually spreads down the age gradient to younger leaves. Ethylene accumulation and signaling throughout the plant causes this age gradient, whereby the transcription factor ORE1 plays a dominant role in rapidly starting the de-greening process preferentially in older leaves. Though ethylene leads to ORE1 protein accumulation independent of age, ORE1 activation via phosphorylation occurs specifically in the older leaves. Such a mechanism ensures ORE1 target activation and senescence only in these older leaves. Although ORE1 protein was already produced in young leaves within one day of submergence, its effects on the transcriptome were minimal during 4 days of submergence. The premature production of ORE1 in young leaves means that during prolonged submergence, when energy levels are low, senescence can be induced without the need to make new ORE1 protein. In such an instance ORE1 only needs to be phosphorylated to induce the transcription of its downstream targets.

ORE1 is arguably one of the best-studied transcription factors controlling leaf senescence in Arabidopsis. Transcription of *ORE1* is, besides EIN3, directly induced by other transcription factors including ATAF1, ATAF2, PIF4, PIF5, ABI5, EEL, PRR9, WRKY71, GI, and ARF2 (Garapati et al., 2015; Nagahage et al., 2018; Sakuraba et al., 2014; Kim et al., 2018; Yu et al., 2021; Kim et al., 2020; Xue et al., 2022). *ORE1* mRNA levels are regulated post transcriptionally by *miR164* (Kim et al., 2009). The low *ORE1* mRNA levels in old leaves under control conditions (Figure 2B & 3B) despite the strong *ORE1* promoter activity (Figure S3B) suggest that *ORE1* mRNA was rapidly broken down, potentially via miR164. ORE1 protein levels are controlled by ubiquitination by the E3 ligase NLA and the E2 conjugase PHO2, and by deubiquitination by the ubiquitin-specific proteases UBP12 and UBP13 (Park et al., 2018, 2019). Lastly, the transactivation activity of ORE1 is activated via its phosphorylation by CPK1 (Durian et al., 2020). The established pathway of miR164-based inhibition of premature ORE1 accumulation was not sufficient to prevent the accumulation of ORE1 in young leaves during flooding. Despite the plethora of regulators that affect the abundance of *ORE1* mRNA and protein, we found that ORE1 abundance does not explain the difference in ORE1 target gene activation between old and young leaves during submergence. Rather, this difference is controlled by post-translational modification of ORE1, which limits its activity to old leaves. ORE1 induces the transcription of its targets via an interaction with the positively charged C-terminus of Mediator complex subunit 19a (MED19A), which recruits RNA Polymerase II to target genes (Cheng et al., 2022). Phosphorylation of a protein typically reduces its charge, and the phosphorylation of ORE1 could potentially facilitate its binding to MED19a. This is also consistent with the impaired transactivation activity of ORE1Δ17, although it still exhibits DNA binding activity (Durian et al., 2020).

Ethylene can freely diffuse across cell membranes and does not require specific transporters to move between cells. The lack of control of ethylene movement requires a plant to have a highly tissue-specific response system to ethylene. This has been described for different cell types (Vaseva et al., 2018; Polko et al., 2011; Cao et al., 1999; Rajhi et al., 2011), but also for similar tissues in different developmental stages (Jing et al., 2005; dela Fuente and Leopold, 1968). Tissue-specific regulation of ethylene responsiveness occurs on many different levels of the ethylene signaling cascade (Stepanova and Alonso, 2009). Since ethylene-insensitive mutants do not induce ORE1-mediated senescence of old leaves during flooding, their old leaves die slower than those of wild-type plants. The young leaves of ethylene-insensitive mutants, on the other hand, die faster than those of wild-type plants. This could be an effect of the impaired ability of ethylene mutants in responding to reactive oxygen species that accumulate during submergence recovery or of other unidentified roles of ethylene in submergence survival (Tsai et al., 2014; Liu et al., 2022). This highlights how ethylene signaling can lead to either the death or survival of a leaf during flooding stress, depending on the age of the leaf.

Our results show that ethylene-induced leaf senescence requires darkness. It is currently unclear if this is an effect of light signaling or of darkness-induced carbon starvation, as both are known to interact with ethylene signaling (Zhong et al., 2012; Shi et al., 2016; Yanagisawa et al., 2003; Kim et al., 2017). In addition to ethylene accumulation, impaired gas diffusion also leads to a decline in oxygen levels in flooded plants. Hypoxia is also considered an important regulatory signal mediating flood survival responses. Hypoxia by itself does not impose a gradient of age-dependent leaf yellowing, and mutants with impaired hypoxia sensing still show age-dependent leaf death during flooding stress (Figure 1D-F). Furthermore, core hypoxia genes are not affected by the loss of *ORE1*, showing that *ORE1* is not upstream of hypoxia signaling (Figure S4D). Based on these results we conclude that the sequential leaf death described here does not appear to involve oxygen sensing and signaling. ORE1 and rice SUB1A are both important regulators of the response to submergence, but both of them are primarily controlled by ethylene rather than hypoxia (Gibbs et al., 2011; Lin et al., 2019). This likely stems from the prevalence of hypoxia in normal plant development and the variation in oxygen concentrations among submerged plant tissues (Weits et al., 2019; Sasidharan et al., 2018).

Ethylene accumulation upon submergence induces senescence of old leaves via the age-dependent phosphorylation of ORE1. Our results suggest that this phosphorylation is independent of CPK1, which is known to phosphorylate ORE1 in vivo (Durian et al., 2020). Future research should focus on how exactly the age-dependent phosphorylation of ORE1 is controlled. Protein kinases and phosphatases themselves are often controlled post-translationally and the interactions between them and their targets can be highly context-specific (Simeunovic et al., 2016; Bhaskara et al., 2019), potentially complicating the identification of the posttranslational regulators of ORE1 during submergence.

The severely reduced diffusion of gases in water means that ethylene will accumulate rapidly in any plant tissue that is completely submerged. This property of ethylene makes it an ideal flood warning cue mediating many flood-adaptive traits (Sasidharan and Voesenek, 2015). However, such high concentrations of ethylene mean that senescence is inevitable for submerged leaves. Therefore, a mechanism that prevents the simultaneous indiscriminate breakdown of all leaf tissue in such a situation is essential to prolong survival. During natural plant ageing, the genetically coordinated process of chlorophyll breakdown during senescence serves to remobilize nutrients towards seed and tuber filling (Yu et al., 2015). The ability to retain chlorophyll has been found to correlate with higher tolerance to submergence and improved post-submergence photosynthesis (Alpuerto et al., 2016; Yeung et al., 2018). For example, the submergence tolerance gene *SUB1A* delays leaf senescence. Like *ORE1*, the expression of *SUB1A* is regulated by ethylene (Fukao et al., 2006). Whereas ORE1 induces chlorophyll degradation, *SUB1A* inhibits it during both submergence and darkness, and thereby contributes to a quiescence strategy during flooding (Xu et al., 2006; Fukao et al., 2006, 2012). However, during prolonged submergence as energy reserves become increasingly limited, senescence would be a beneficial option. In such a situation, a sequential dismantling of older leaves would make available energy and nutrient reserves that can be redirected to sustain younger leaves and the meristem. This sacrificial use of older leaves would serve to enhance growth and photosynthesis recovery when floodwaters subside. Understanding how plants coordinate which tissues are broken down under stressful conditions could help in developing more stress-tolerant crop varieties, as the role of NAC domain transcription factors in senescence is conserved across many plant species (Podzimska-Sroka et al., 2015).

## Methods

### Plant material

*ore1-1* (SALK_090154): described in (He et al., 2005). Ordered from NASC

*ore1-2* (SAIL_694_C04): described in (Kim et al., 2020). Ordered from NASC

*cpk1-1* (SALK_096452): described in (Durian et al., 2020). Ordered from NASC

*35S:ORE1:* described in (Matallana-Ramirez et al., 2013). Gift from Salma Balazadeh

*35S:ORE1Δ17:* described in (Durian et al., 2020). Gift from Salma Balazadeh

*ein2-5*: described in (Alonso et al., 1999). Ordered from NASC

*ein3eil1*: described in (Alonso et al., 2003). Ordered from NASC

*pco124*: described in (Masson et al., 2019). Gift from Daan Weits

*erfVII*: described in (Abbas et al., 2015). Gift from Daan Weits

*prt6-1*: described in (Garzón et al., 2007). Gift from Angelika Mustroph

*35S:EIN3-GFP ein3eil1*: described in (Xie et al., 2015). Gift from Shi Xiao

RPS5a::XVE>>ORE1-GFP: Described in (Gao et al., 2018). Gift from Moritz Nowack

*pORE1:ORE1-HA ore1-1*: this study

*pORE1:GUS*: this study

All *Arabidopsis thaliana* (Arabidopsis) lines were in the ecotype Col-0 (Columbia-0) background

### Generating transgenic lines

Genomic DNA from an Arabidopsis ecotype Col-0 leaf was extracted using phenol:chloroform:isoamyl alcohol. The *ORE1* genomic region, including introns, 5’UTR, and a 1624bp promoter, was amplified from this DNA using primers 5383 & 5384 (Supplemental table 2) and inserted into the pJET1.2 vector (Thermo Fisher, K1231) according to the manufacturer’s instruction. For the *pORE1:ORE1-HA* line the entire fragment without the stop codon was amplified from this vector using primers 5383 & 5510, and the HA tag was added using primers 5383 & 5783. For the *pORE1:GUS* line the *ORE1* promoter was amplified using the primers 5383 & 5712. Adapters for binary LIC vectors pPLV01 and pPLV13 (De Rybel et al., 2011) were added to the *pORE1:ORE1-HA* and *pORE1* fragments using primers 5804 & 5761 and 5739 & 5740, respectively. The fragments were inserted into their respective vectors via ligation-independent cloning as described in (De Rybel et al., 2011). These vectors were introduced into *Agrobacterium tumefaciens* strain AGL-1 via electroporation, and *ore1-1* and Col-0 Arabidopsis plants were transformed using the floral dip method (Logemann et al., 2006). Independent T_1_ transformants were selected on plates containing 50µM Basta/PPT, homozygous T_3_ or T_4_ lines were used in all experiments.

### Plant growth & treatments

Seeds were sowed on Primasta soil mix and stratified in the dark for 3 to 4 days before being transferred to a short-day condition climate chamber (20C, 9h light 15h dark, 70%RH, ∼140-180PAR either LED or fluorescent light). After germinating for nine days, seedlings were transplanted to individual (5.5cm diameter, 5cm height) pots consisting of 2:1 perlite:soil mix, which were covered with a black mesh to prevent soil from floating out of the pot during submergence. 1 liter of 0.5x Hoagland medium was added to each tray of 42 pots. When plants reached the 10-leaf stage they were submerged in complete darkness at 20C for the indicated amount of time, and were left to recover for the indicated amount of time in the original climate chamber. Submergence treatment for the timelapse videos (Supplemental videos 1 & 2) was done at 1 PAR, pictures were taken every 30 minutes over two weeks using a Nikon D750 camera. Ethylene treatments were done in 22.5l desiccators as described in Hartman et al. (2019). Hypoxia treatments were done by mixing N_2_ and air to a concentration of 5% O_2_, which was flushed through a desiccator for 1h. Desiccator valves were then closed and 5-10ppm ethylene was injected with a syringe. For leaf death quantification, leaves were scored as dead when more than half of the leaf area had desiccated after 3 days of post-submergence recovery in the light.

### Green and senescing leaf area quantification

Pictures of Col-0 and *ore1-1* plants were taken with a Nokia 8 phone camera. Individual pixels in each picture were classified into either “green”, “senescing”, “dead”, or “background” using the “Naïve Bayes Multiclass” module within PlantCV (Fahlgren et al., 2015). ImageJ was used to count the amount of pixels in the green and senescing categories, and this was plotted relative to the amount of green pixels before the start of treatment for Figure S2H. For Figure 2H the amount of green and senescing pixels of the same plants were added together and converted into an area in cm^2^.

### Seed yield

For seed yield measurements, plants were either kept in short-day control conditions or submerged for 6 days in darkness and then returned to control conditions. Watering was stopped once the first siliques started to dry out and plants were left to dry out until all siliques had ripened.

### Chlorophyll quantification

For chlorophyll measurements, individual old or young leaf blades of the indicated genotypes were cut off and put into 1.5ml Eppendorf tubes containing 1ml DMSO at the indicated timepoints. Tubes were incubated in a shaking water bath at 60C for 30 minutes in darkness, and were then left to cool down to room temperature for another 30 minutes in darkness. 200µl of each DMSO solution was pipetted into a 96-well plate and absorption was measured at 647nm, 664nm, and 750nm using a spectrophotometer plate reader (Synergy HT Multi-Detection Microplate Reader; BioTek Instruments). Chlorophyll A was calculated as 13.71*(664nm-750nm)-2.858*(647nm-750nm), chlorophyll B was calculated as 22.39*(647nm-750nm)-5.42*(664nm-750nm). Leaves were dried at 80C for 48h before measuring their dry weight on a Mettler-Toledo MX5 microbalance. Total chlorophyll was calculated by adding chlorophyll A and B together and dividing them by the measured dry weight.

### Ion leakage

Five leaves per replicate of indicated tissues were pooled together in a 15ml tube containing 3ml distilled water, and were gently shaken for 3 hours. The concentration of ions in the solution was measured using a Horiba EC-33 conductivity meter. Plant tissue was then boiled for 20 minutes to destroy all membranes, and ion leakage was measured again to determine the total ion content. Relative ion leakage was calculated as the proportion of the conductivity before boiling to the conductivity after boiling.

### Gene expression

RNA was extracted from the indicated tissues using the Qiagen plant RNeasy mini kit, including an on-column DNAse treatment, according to the manufacturer’s instructions. qPCR data shown in figures 3B and S3B was done using the Spectrum RNA extraction kit (Sigma-Aldrich), followed by DNase treatment using AMPD1 DNase I (Sigma-Aldrich) to ensure that *miR164b* would not be excluded by the size exclusion limit of the Qiagen kit.

Extracted RNA was converted into cDNA using RevertAid H Minus Reverse Transcriptase (Thermo Scientific). For qPCR 20ng of cDNA was used per 5µl reaction, using SYBR Green master mix (BioRad) and the primers indicated in Supplemental table 1.

### GUS staining

Whole rosettes of 10-leaf *pORE1:GUS* plants were cut off at the indicated timepoints and were fixated in 90% acetone for 20 minutes. Plants were then washed twice for 10 minutes in GUS washing buffer (0.1M phosphate buffer pH=7, 10mM EDTA, 2mM K_3_Fe(CN)_6_) under vacuum. Plants were then stained with GUS washing solution (0.1M phosphate buffer pH=7, 10mM EDTA, 1mM K_3_Fe(CN)_6_, 1mM K_4_Fe(CN)_6_*3H2O, 0.5mg/ml X-Gluc) for 10 minutes under vacuum, followed by 20 hours at 37C. Staining was stopped by incubating the plants with 3:1 acetic acid:ethanol for 1h, and were cleared by washing with 70% ethanol. Plants were scanned using an Epson V800 scanner.

### RNA sequencing

Between 8 and 16 young and old leaves of Col-0 and *ore1-1* were harvested before submergence, after 4 days of dark submergence, and after 6 hours of post-submergence recovery in the light. Additional Col-0 samples were harvested after 2 days (old and young leaves) and six days of submergence (young leaves only), and after 1, 3, and 24 hours of recovery (old and young leaves). RNA was extracted using the Qiagen RNeasy Plant Mini kit, genomic DNA was removed by treating the samples with AMPD1 DNase I (Sigma-Aldrich). Libraries were constructed by Macrogen using the TruSeq Stranded mRNA LT Sample Prep Kit (Illumina). Libraries were sequenced on an Illumina NovaSeq6000 platform NovaSeq6000 via paired-end sequencing of 150bp reads. Sequenced libraries were trimmed of adapter sequences using FastQC (Babraham Bioinformatics). Cleaned reads were aligned to the Araport11 transcriptome using Kallisto (Bray et al., 2016). Genes were determined as differentially expressed when FDR<0.05 and |log_2_FC|>1, as calculated using R packages EdgeR and limma (Supplemental table 3). Fold changes and p-values for all timepoints were also calculated compared to non-submerged old leaves of Col-0, these were used in Figures S4D and S6A and can be found in Supplemental table 4.

### ORE1 binding site density

To determine the density of ORE1 binding sites in the promoters of putative target genes, genes were selected from the RNAseq dataset that are induced significantly stronger in Col-0 old leaves than *ore1-* 1 old leaves after 4d of dark submergence. Promoters (1kb upstream and 100bp downstream of the transcriptional start site) of these 287 genes were extracted from the TAIR9 genome sequence using the GenomicRanges R package (Lawrence et al., 2013). These promoters were scanned for the occurrences of the ORE1 motifs VMGTR_N5-6_YACR and TDRCGTRHD, allowing one mismatch (Matallana-Ramirez et al., 2013; Olsen et al., 2005). The density of motif centers along the promoter sequences was corrected for the number of scanned promoters and plotted using ggplot2.

### Electrophoretic mobility shift assay

EMSAs were performed as previously described by (Wu et al., 2012). ORE1-GST protein was purified as previously described by (Durian et al., 2020). Binding reactions were performed using the Odyssey infrared EMSA kit (LI-COR) following the manufacturer’s instructions. DNA-protein complexes were separated in a 6% (w/v) retardation gel (EC6365BOX, Invitrogen). DY682 signal was detected using the Odyssey infrared imaging system from LI-COR.

### ChIP-qPCR

For ChIP, 10-leaf stage Col-0 and *pORE1:ORE1-HA ore1-1* plants were submerged for one day in darkness to induce ORE1-HA protein accumulation. Chromatin was extracted from 1.5g of whole-rosette tissue for each replicate. Protein-DNA complexes were immunoprecipitated using anti-HA antibodies (Miltenyi Biotec) (Kaufmann et al., 2010). After reversion of the cross-linking, DNA was purified with the QIAquick PCR Purification Kit (Qiagen) and was analyzed by qPCR. Enrichment of ORE1 at the target promoters was calculated relative to Col-0, significance was determined by comparing the enrichment at each of the target loci to that of the negative control (AT2G22180).

### Western blotting

Five leaves of the indicated age were pooled together after the indicated treatment and frozen in liquid nitrogen. Protein was extracted using RIPA buffer (Hartman et al., 2019) and quantified using a Pierce BCA kit. 20-50µg of protein was loaded onto a stain-free 4-15% gel (BioRad), the Rubisco large subunit was visualized using stain-free gel imaging. Proteins were transferred from the gel to a 0.2µm PVDF membrane using a Bio-Rad trans-blot system for 7 minutes, efficient transfer was verified by imaging the stain-free blot afterwards. The blot was blocked overnight at 4C in TBS-T + 5% milk. Primary antibody (1:1000, anti-GFP (Roche, #11814460001) or anti-HA-HRP (Thermo Fisher, 26183-HRP)) was incubated for 1h @ RT. The blot was washed 4 times 10 minutes with TBS-T. In the case of anti-GFP blots, the membrane was incubated with a secondary antibody (1:2500 rabbit anti-mouse, Cell signaling #7076) for 1h at RT and the membrane was washed 3x with TBS-T and 2x with TBS, 5 minutes each. The membrane was incubated with Femto (Thermo Fisher) and imaged under a ChemiDoc imaging system (BioRad) to visualize HRP activity.

To identify phosphorylated proteins, 50µg of protein extract was precipitated by incubating the sample with 4x the volume of the protein sample 100% ice-cold acetone for 1h at −20C. After precipitation the samples were centrifuged for 10 minutes at 13,000g at 4C and the supernatant was removed. The pellet of precipitated proteins was resuspended in 12µl water and 3µl 5x sample loading buffer (250mM Tris (pH 6.8) 25% glycerol, 10% SDS, 0.05% bromophenol blue) containing 5% beta-mercaptoethanol. Samples were boiled for 5 minutes at 95C to denature the proteins and were separated via electrophoresis on a SuperSep Phos-Tag 7.5% gel with 50µmol/l Phos-Tag (198-17981, Fujifilm Wako, Japan). The gel was run at a stable 20mA for 2.5h. After electrophoresis, the gel was washed twice in running buffer containing 10mM EDTA for 10 minutes and once in running buffer without EDTA. Transferring the proteins, blocking the membrane, antibody incubation, and imaging were done identically to non-phostag ORE1-HA Western blots. The large subunit of rubisco was imaged after staining for ORE1-HA using 0.1% (w/v) Ponceau S to verify equal loading.

### Estradiol treatment

Entire *RPS5a::XVE>>ORE1-GFP* plants were sprayed twice daily with 100µM estradiol (from a 20mM estradiol stock in ethanol) in water or a mock solution (0.5% ethanol). Leaves 1, 3, and 7 were harvested after 8 days and snap-frozen in liquid nitrogen.

### Statistical analysis

All statistical tests were done in R version 3.6.1 using the indicated statistical tests, differences were deemed significant at p<0.05.

## Author contributions

Conceptualization: TR, HvV, RS; Investigation: TR, HvV, MS, C-YL, MBD; Data analysis: TR, HvV, MS; Funding acquisition: RS; Methodology: TR, HvV, MS, C-YL; Supervision: SB, RS; Writing – original draft: TR, RS; Writing – review & editing: TR, HvV, SB, RS.

## Acknowledgments

We would like to thank Bernhard Würzinger and Markus Teige for their input on PhosTag Western blots and Yorrit van de Kaa for harvesting seeds. This work was financially supported by the Netherlands Organization for Scientific Research grant 016.VIDI.171.006 to TR and RS and grant ALWOP.419 to HvV. SB thanks the Max Planck Institute of Molecular Plant Physiology (MPIMP) and Leiden University for funding.

## Data and code availability

RNAseq data has been deposited at the European Nucleotide Archive under accession number PRJEB57289. Transcript abundance in the RNAseq data can also be explored in a Shiny app at https://utrecht-university.shinyapps.io/Rankenberg2022/. All other data and code are available from the lead contact upon request.

