## Supplementary material for "Differential leaf flooding resilience in *Arabidopsis thaliana* is controlled by age-dependent ORESARA1 activity": SI data

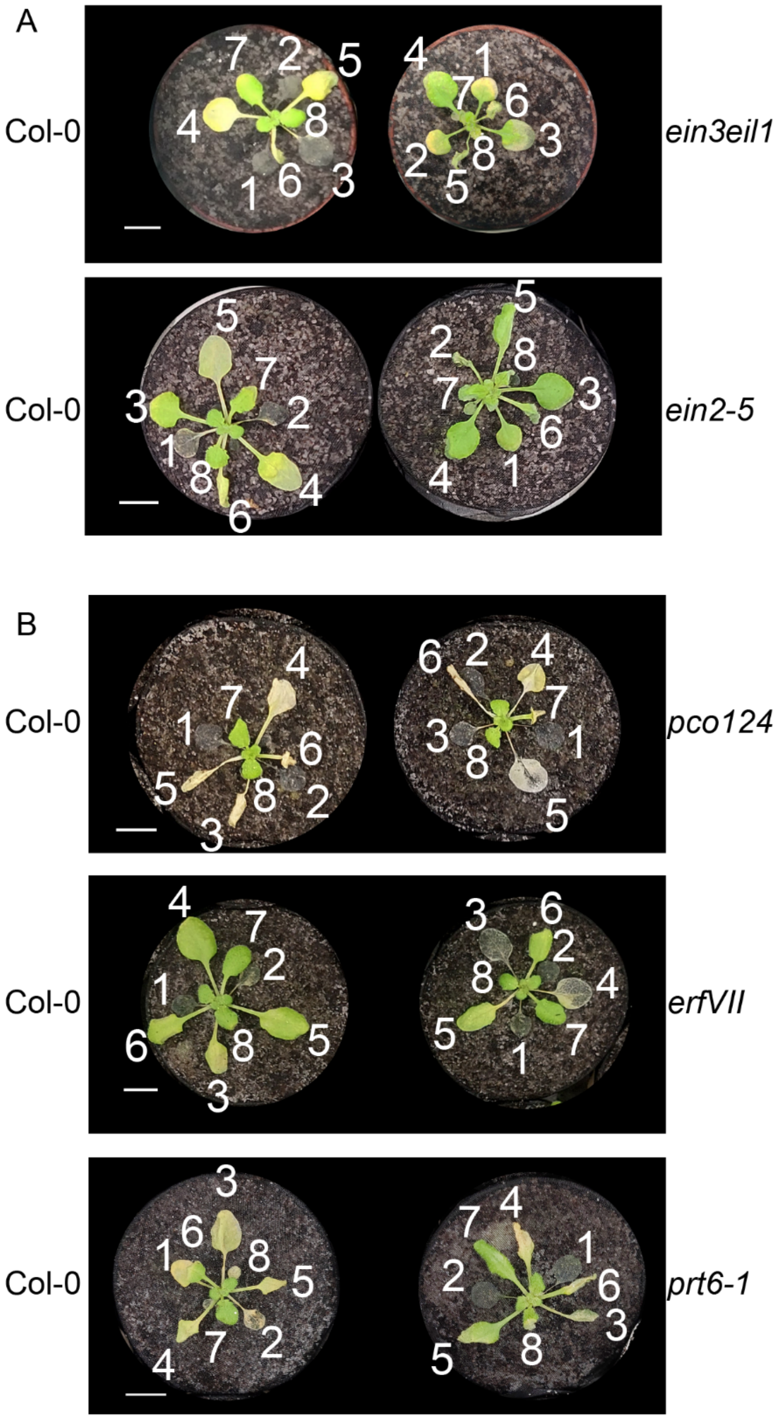


**Figure S1**

A) Representative images of *ein3eil1* and *ein2-5* mutants after 5 (*ein2-5*/Col-0) or 8 (*ein3eil1*/Col-0) days of submergence followed by one day of recovery

B) Representative images of *pco124*, *erfVII*, and *prt6-1* mutants after 5 (Col-0/*pco124*) or 4 (Col-0/*erfVII*, Col-0/*prt6-1*) days of submergence followed by one day of recovery.

Numbers indicate leaf numbers based on the order of emergence. Scale bars indicate 1cm

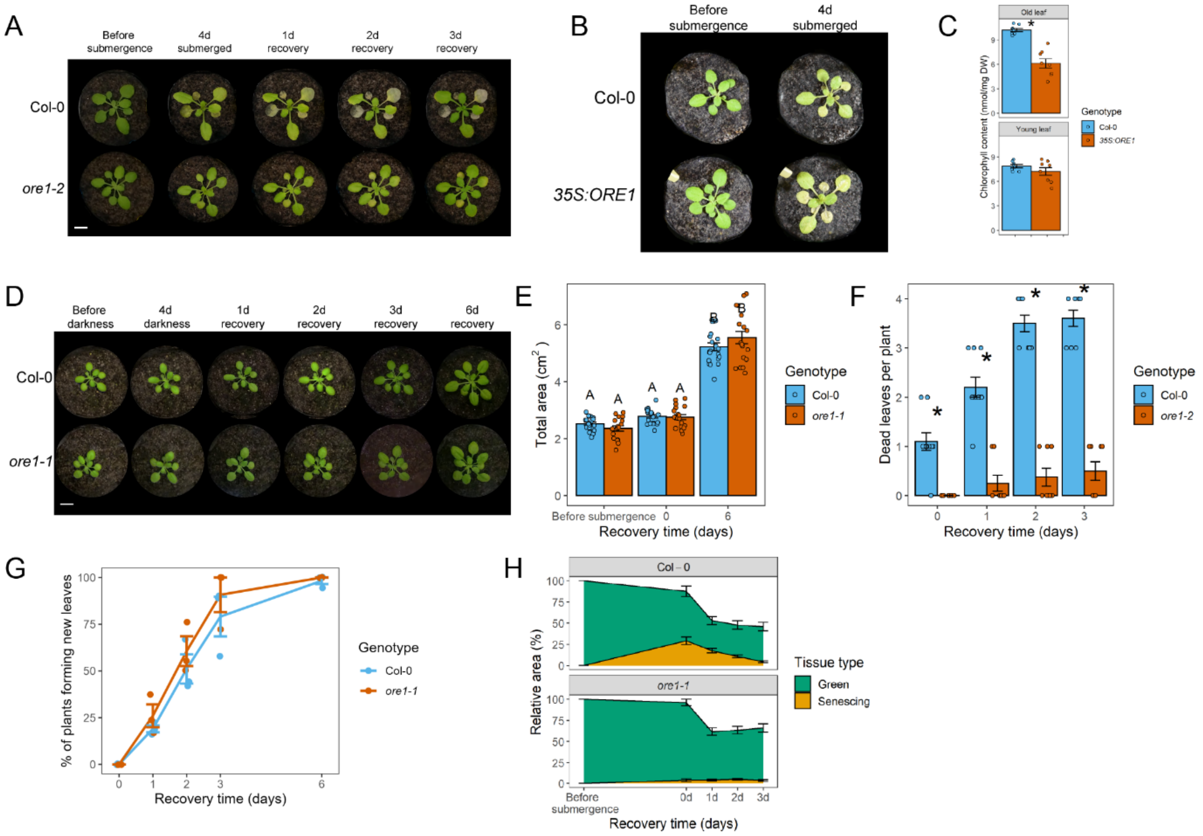


**Figure S2**

A) *ore1-2* mutants show reduced yellowing of old leaves after recovery from four days of submergence. Representative images of Col-0 and *ore1-2* plants at the indicated timepoints.
B) *35S:ORE1* plants show increased leaf senescence after 4 days of submergence. Representative images of Col-0 and *35S:ORE1* plants at the indicated timepoints.
C) Chlorophyll content of old and young leaves of Col-0 and *35S:ORE1* plants after 4 days of submergence. Asterisks indicate signficant differences (t-test)
D) Four days of darkness does not induce leaf senescence in Col-0 or *ore1-1*. Representative images of Col-0 and *ore1-1* plants at the indicated timepoints.
E) Four days of darkness does not affect the rosette area of Arabidopsis plants in an *ORE1*-dependent manner. Different letters indicate significant differences between groups (two-way ANOVA + Tukey's post-hoc test). n=16-20 per genotype.
F) Dead leaves of Col-0 and *ore1-2* during recovery from 5 days of submergence. Asterisks indicate signficant differences (t-test), n=8-10.
G) The rate of new leaf formation while recovering from 5 days of submergence is not significantly different between Col-0 and *ore1-1* plants (t-test). Dots represent three independent experiments, consisting of 8-21 plants per genotype each.
H) Relative green and senescing areas of Col-0 and *ore1-1* before and after 4 days of submergence, and at the indicated recovery timepoints, as quantified by PlantCV. n =17 (Col-0), 20 (*ore1-1*).
Scale bars indicate 1cm

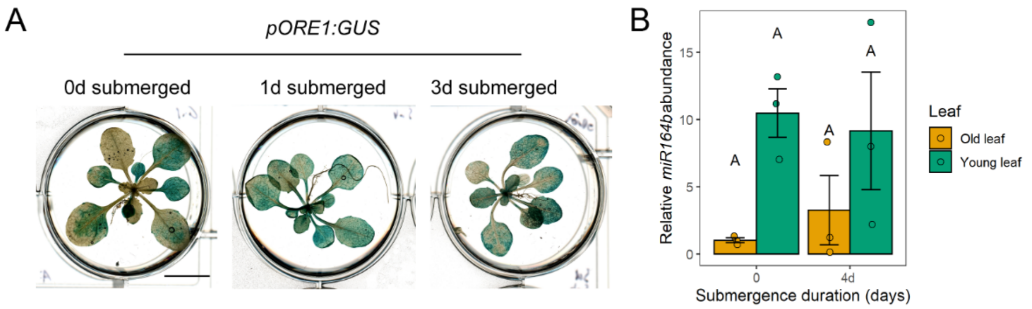


**Figure S3**
A) GUS staining of *pORE1:GUS* shows that *ORE1* promoter activity is limited to old leaves and cotyledons in non-submerged plants but is induced in all leaves upon submergence. Scale bar indicates 1cm.
B) *miR164b* pri-miRNA abundance in old and young leaves before and after four days of submergence. Different letters indicate significant differences between groups (two-way ANOVA + Tukey's post-hoc test). N=4 for non-submerged plants, n=3 for submerged plants.

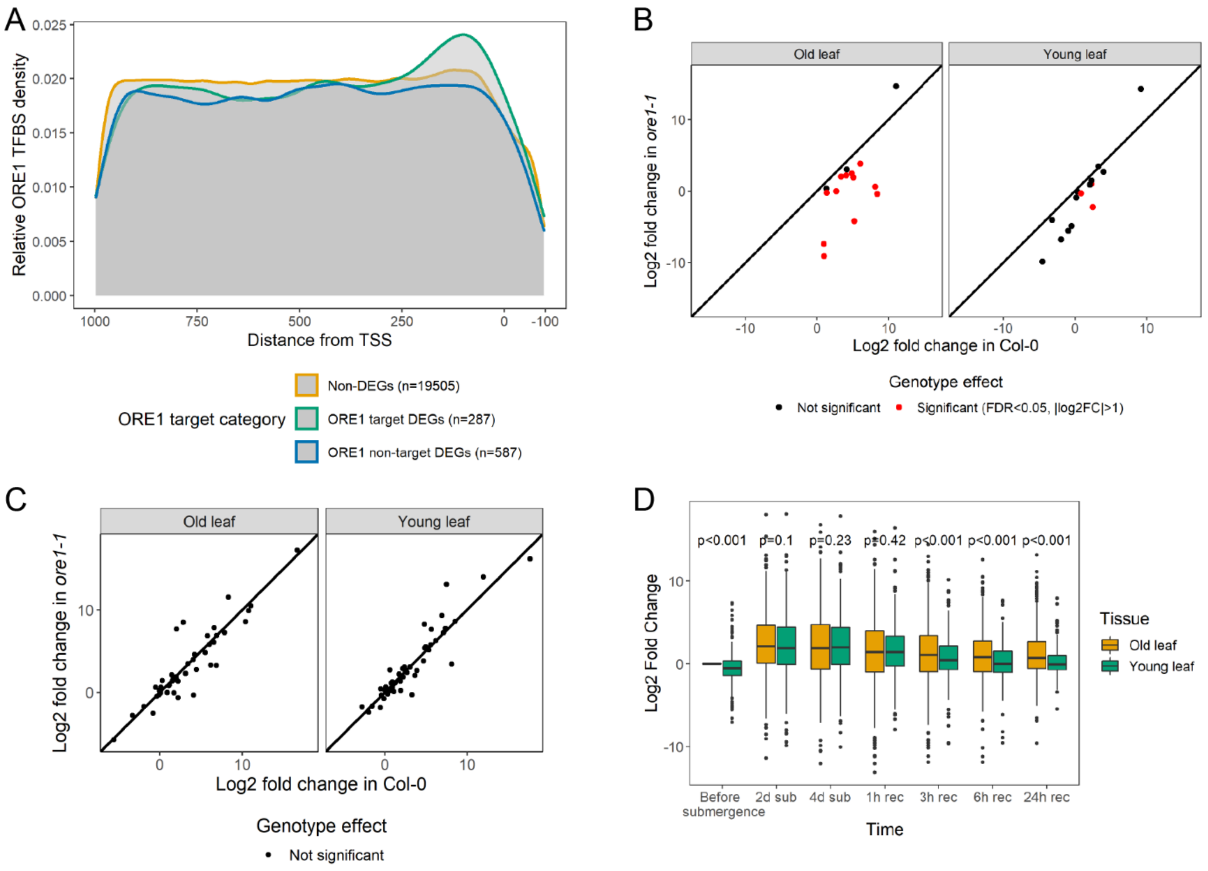


**Figure S4**
A) Density of *ORE1* binding sites (TDRCGTRHD/VMGTRN5-6YACR, Olsen et al., 2005; Matallana-Ramirez et al., 2013) in the promoters of Non-DEGs, DEGs with lower expression in *ore1-1* than Col-0 in old leaves (ORE1 target DEGs), and all other DEGs between Col-0 and *ore1-1* in old leaves.
B) *In vivo* confirmed ORE1 target genes (n=15, supplemental table S3) show more differences between Col-0 and *ore1-1* in their response to four days of submergence in old leaves than in young leaves.
C) The response of none of the core hypoxia genes (Mustroph et al., 2009) to four days of submergence is significantly different between Col-0 and *ore1-1*.
D) Expression of EIN3 target genes (Chang et al., 2013) is similar in old and young leaves during submergence but is higher in old leaves before and after submergence (Wilcoxon Rank Sum test).

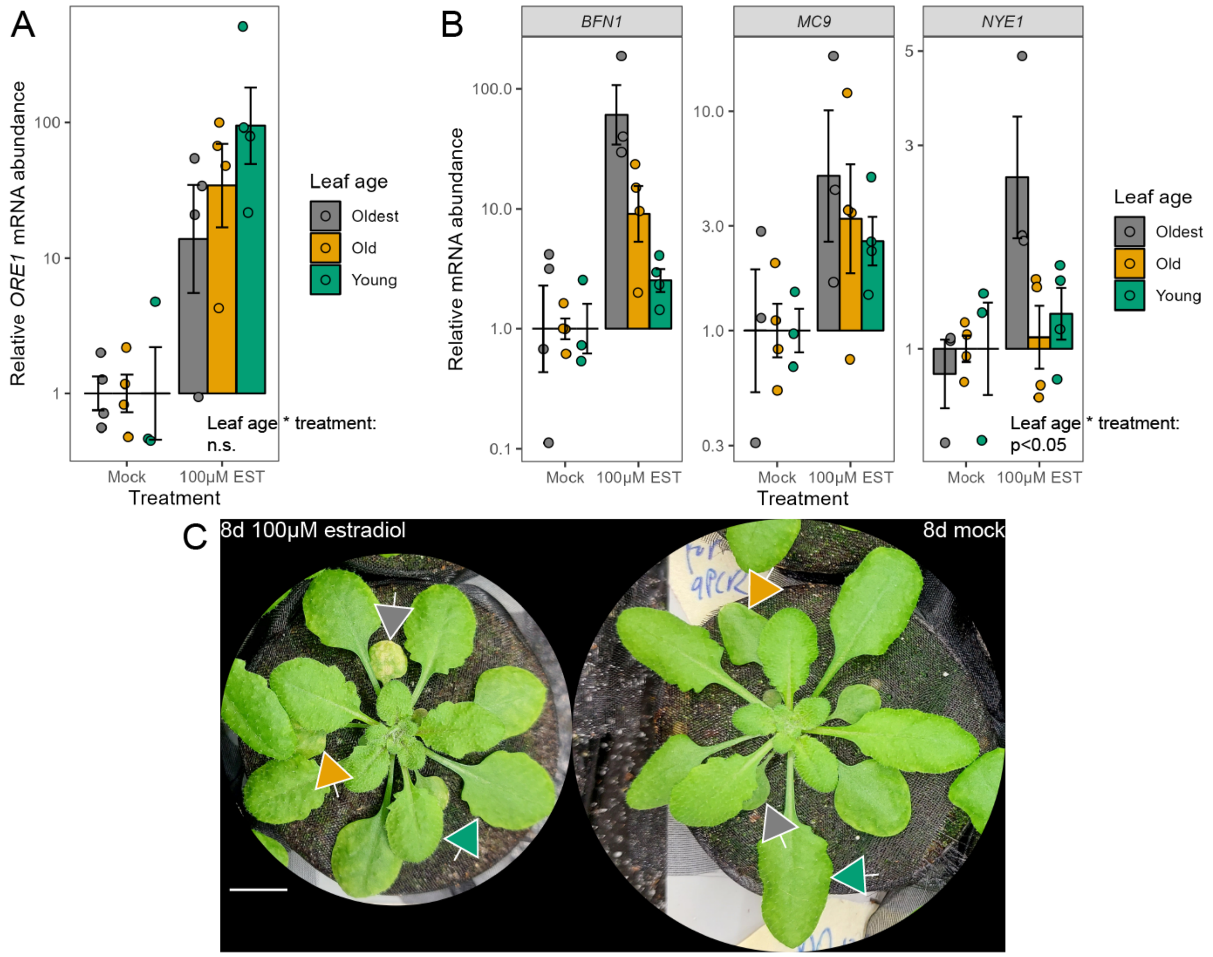


**Figure S5**

A) mRNA abundance of *ORE1* in the oldest leaf (number 1), an old leaf (leaf 3), and a young leaf (leaf 7) of *RPS5a::XVE>>ORE1-GFP* plants after 8 days of estradiol or mock treatment. n=3-4 per sample, each consisting of 2 leaves of different plants pooled together. The indicated p-value is the interaction effect in a two-way ANOVA for the effect of leaf age and the treatment on *ORE1* expression, with the harvested plant as a blocking effect.

B) mRNA abundance of ORE1 targets *BFN1*, *MC9*, and *NYE1* in the oldest leaf (number 1), an old leaf (leaf 3), and a young leaf (leaf 7) *RPS5a::XVE>>ORE1-GFP* plants after 8 days of estradiol or mock treatment. n=3-4 per sample, each consisting of 2 leaves of different plants pooled together. The indicated p-value is the interaction effect in a two-way ANOVA for the effect of leaf age and the treatment on the expression of ORE1 target genes, with the harvested plant and the specific ORE1 target gene as a blocking effect.

C) Phenotype of *RPS5a::XVE>>ORE1-GFP* plants after 8 days of estradiol or mock treatment. Arrowheads indicate the leaves harvested in panels A and B, colors correspond to the colors in the legends of these panels. Scale bar indicates 1cm.

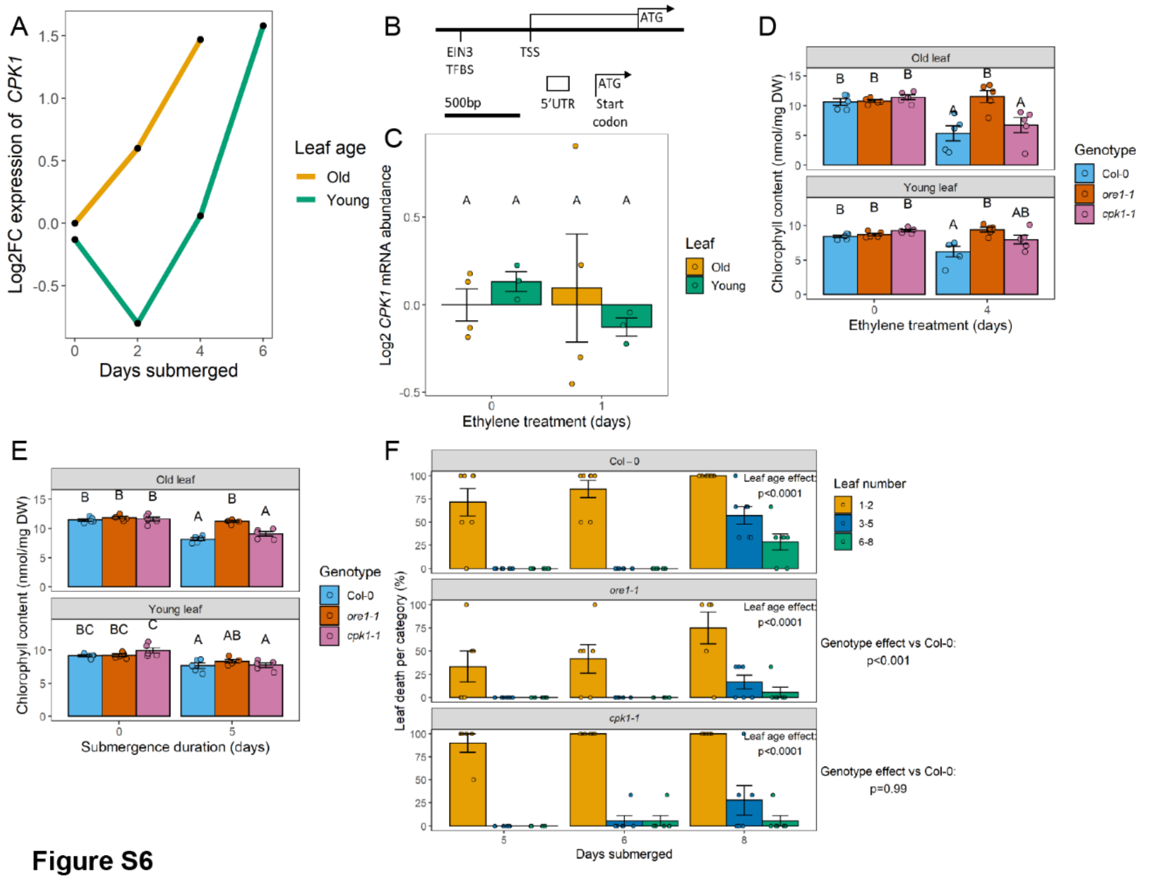


**Figure S6**

A) Expression of *CPK1* during submergence in old and young leaves, based on RNAseq data. Expression is calculated relative to that of non-submerged old leaves.

B) Schematic overview of the EIN3 binding site (AYGWAYCT) in the promoter of *CPK1*, 443bp from its transcriptional start site

C) Expression of *CPK1* in response to ethylene in darkness, as determined by qPCR. Different letters indicate significant differences between groups (two-way ANOVA + Tukey's post-hoc test). N=4 for old leaves, n=3 for young leaves. Two leaves were pooled together per sample.

D-E) Chlorophyll content of Col-0, *ore1-1*, and *cpk1-1* plants before and after 5 days of submergence treatment in darkness (D) or 4 days of ethylene treatment in darkness (E). Different letters indicate significant differences between groups (two-way ANOVA + Tukey's post-hoc test), n=5 per sample.

F) Col-0, *ore1-1*, and *cpk1-1* all show age-dependent leaf death, this process is slowed down in *ore1-1* but not in *cpk1-1*. The leaf age effect was determined by two-way ANOVA (leaf age * time). The genotype effect was determined by two-way ANOVA (genotype + leaf age) followed by Tukey's post-hoc test, p-values correspond to the comparison between either mutant and Col-0.

Error bars indicate SEM.
